## Supplemental Data for "Hedgehog signaling via its ligand DHH acts as cell fate determinant during skeletal muscle regeneration"

### Supplemental Figures and Table

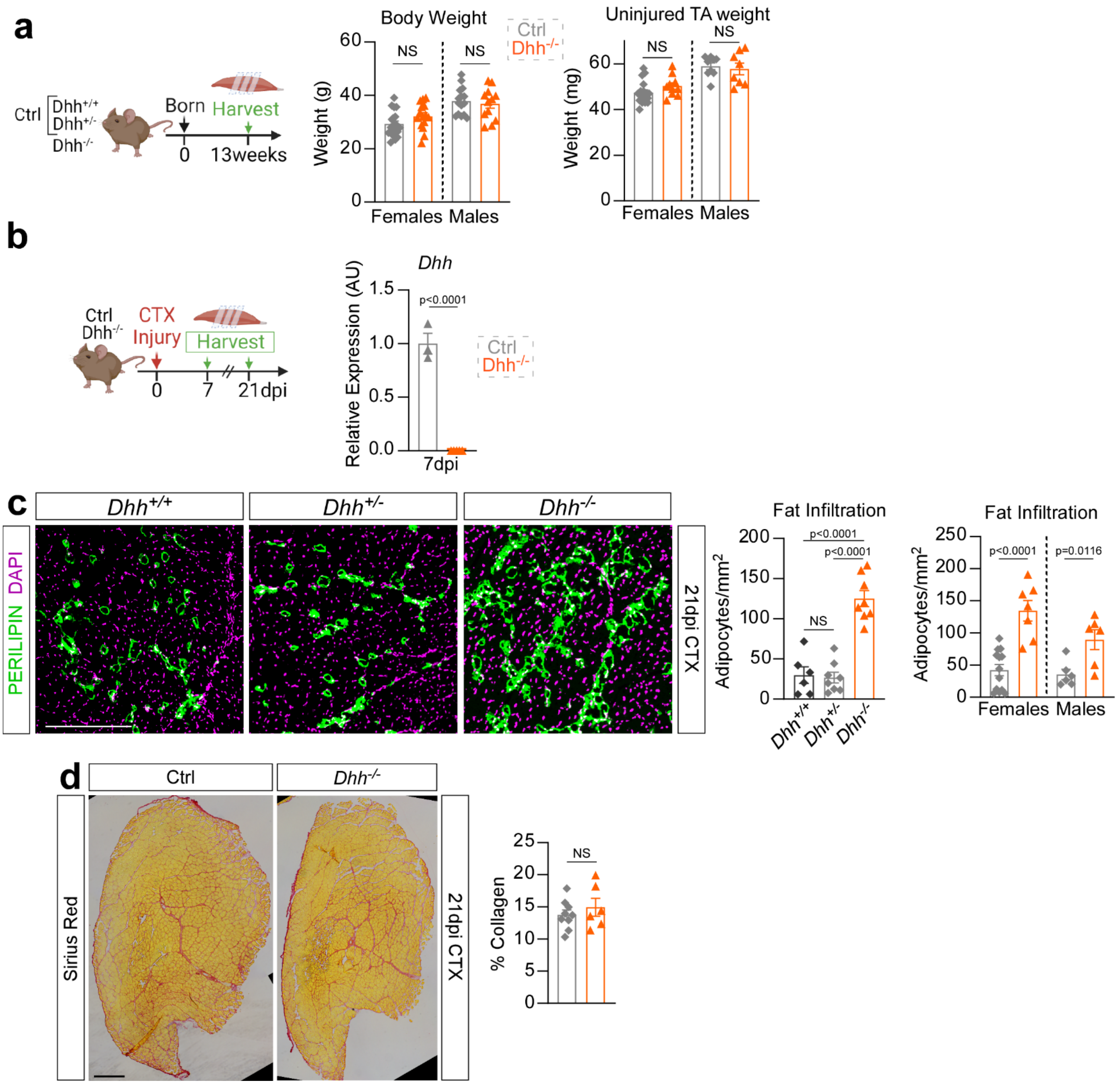


**Supplemental Figure 1. Validation of the *Dhh*$^{-/-}$ mouse model.**

**a)** Experimental outline. Body weight (g) of 13-week-old *Dhh*$^{-/-}$ (females, n=15 mice; males, n=12 mice) and ctrl mice (females, n=19 mice; males n=15 mice). Wet-weight of uninjured TAs (mg) of *Dhh*$^{-/-}$ (females, n=10 TAs; males, n=8 TAs) and ctrl mice (females, n=17 TAs; males n=9 TAs). **b)** Experimental outline. RT-qPCR of *Dhh* expression 7 days after CTX injury of *Dhh*$^{-/-}$ (n=5 TAs) and ctrl (n=3 TAs) mice. **c)** *Left*: Immunofluorescence of adipocytes (PERILIPIN$^{+}$, green) 21 days post CTX injury of *Dhh*$^{+/+}$, *Dhh*$^{+/-}$ and *Dhh*$^{-/-}$ mice. Nuclei were visualized with DAPI (purple). Scale bars: 250 µm. *Right:* Quantification of adipocytes per injured area (mm$^2$) 21 days post CTX injury in *Dhh*$^{+/+}$ (n=6 TAs), *Dhh*$^{+/-}$ (n=8 TAs) and *Dhh*$^{-/-}$ (n=8 TAs) mice. Adipocyte quantification of *Dhh*$^{-/-}$ and ctrl mice separated by females (*Dhh*$^{-/-}$ n=7 TAs; ctrl n=13 TAs) and males (*Dhh*$^{-/-}$ n=6 TAs; ctrl n=6 TAs). **d)** *Left:* Histological Sirius red staining 21 days post CTX injury in *Dhh*$^{-/-}$ and ctrl mice. Scale bars: 500 µm. *Right:* Quantification of percent of collagen (red) over total TA area 21 dpi in *Dhh*$^{-/-}$ (n=6 TAs) and ctrl mice (n=9 TAs). All data are represented as mean ± SEM. An unpaired two-tailed t test or a one-way ANOVA followed by a Dunnet's multiple comparison was used.

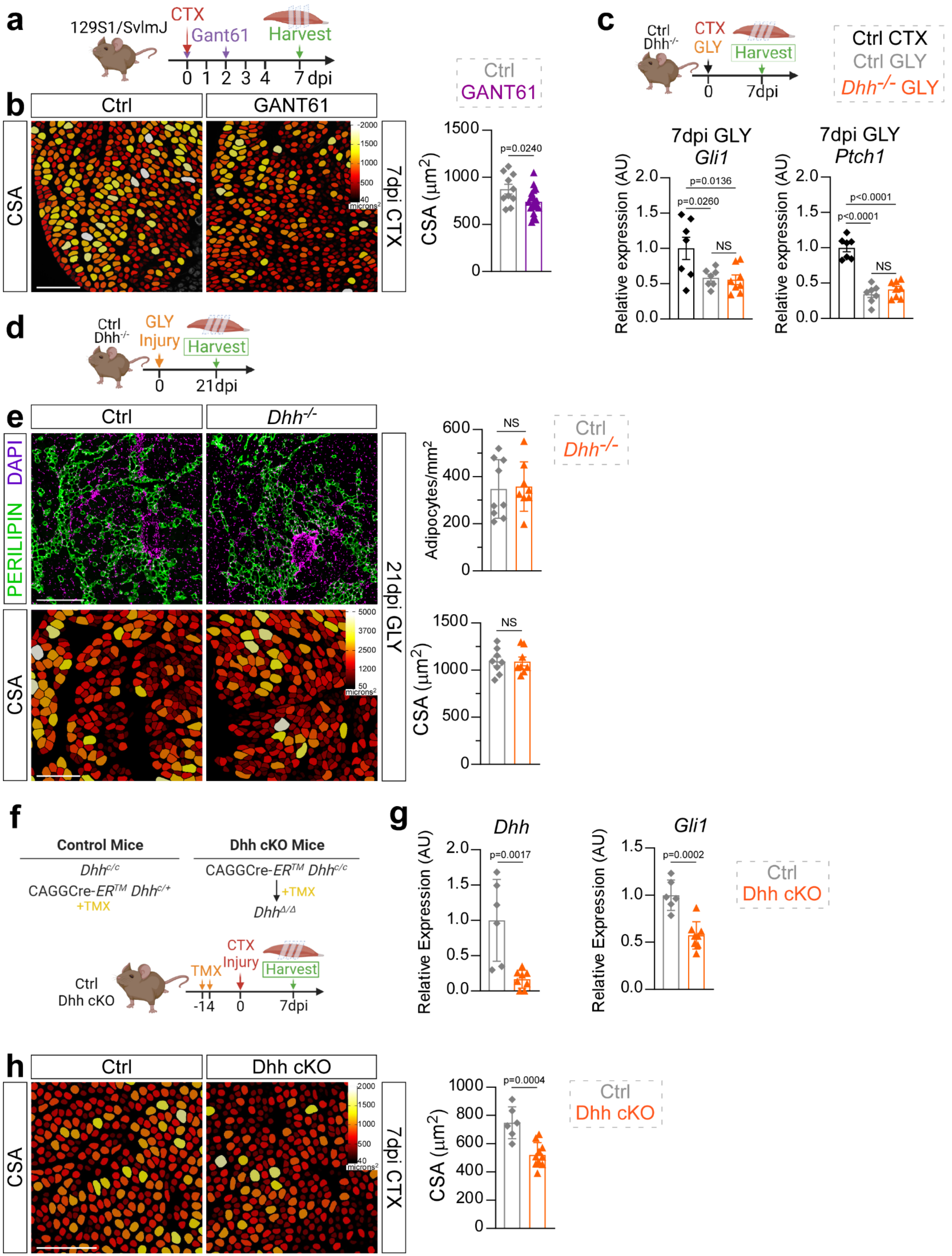

a
129S1/SvlmJ
CTX
Gant61
Harvest
0 1 2 3 4 7 dpi
b
Ctrl
GANT61
CSA
7dpi CTX
2000
1500
1000
500
40
microns²
Ctrl
GANT61
CSA (μm²)
1500
1000
500
0
p=0.0240
c
Ctrl
Dhh-/-
CTX
GLY
Harvest
0
7dpi
Ctrl CTX
Ctrl GLY
Dhh-/- GLY
7dpi GLY
Gli1
Relative expression (AU)
2.0
1.5
1.0
0.5
0.0
p=0.0136
p=0.0260
NS
7dpi GLY
Ptch1
Relative expression (AU)
p<0.0001
p<0.0001
NS
d
Ctrl
Dhh-/-
GLY
Injury
Harvest
0
21dpi
e
Ctrl
Dhh-/-
PERILIPIN DAPI
CSA
21dpi GLY
5000
3700
2500
1200
50
microns²
Adipocytes/mm²
600
400
200
0
NS
Ctrl
Dhh-/-
CSA (μm²)
1500
1000
500
0
NS
f
Control Mice
Dhh cKO Mice
Dhhc/c
CAGGCre-ERTM Dhhc/+
+TMX
CAGGCre-ERTM Dhhc/c
+TMX
DhhΔ/Δ
Ctrl
Dhh cKO
TMX
CTX
Injury
Harvest
-14
0
7dpi
g
Dhh
Relative Expression (AU)
2.0
1.5
1.0
0.5
0.0
p=0.0017
Gli1
Relative Expression (AU)
1.5
1.0
0.5
0.0
p=0.0002
Ctrl
Dhh cKO
h
Ctrl
Dhh cKO
CSA
7dpi CTX
2000
1500
1000
500
40
microns²
CSA (μm²)
1000
800
600
400
200
0
p=0.0004
Ctrl
Dhh cKO

**Supplemental Figure 2. Validation of DHH loss of function data and Dhh cKO mouse model.**
**a)** Experimental outline. **b)** *Left:* Myofibers color-coded according to size (μm$^2$) of Gant61- and vehicle treated mice 7 days post CTX injury. Scale bar: 250 μm. *Right*: Average CSA (μm$^2$) of vehicle (n=10 TAs) and Gant61 (n=19 TAs) treated mice 7 days post CTX. **c)** *Top*: Experimental outline. *Bottom*: RT-qPCR of expression of *Gli1* and *Ptch1* 7 days post CTX in ctrl (n=7 TAs); and 7 days post GLY injury in ctrl (n=7 TAs) and *Dhh*$^{-/-}$ (n=8 TAs) mice. **d)** Experimental outline. **e)** *Left*: Immunofluorescence of adipocytes (PERILIPIN$^+$, green) and nuclei visualized with DAPI (purple); color-coded myofibers according to CSA of *Dhh*$^{-/-}$ and ctrl mice 21 days post GLY. Scale bars: 250 μm. *Right*: Quantification of adipocytes per injured area (mm$^2$) 21 days post GLY in *Dhh*$^{-/-}$ (n=8 TAs) and ctrl (n=9 TAs) mice. Average CSA (μm$^2$) of *Dhh*$^{-/-}$ (n=8 TAs) and ctrl (n=8 TAs) mice 21 days after GLY injury. **f)** Experimental outline. **g)** RT-qPCR of expression of *Dhh* and *Gli1* 7 days post CTX in Dhh cKO (n=8 TAs) and ctrl (n=6 TAs) mice. **h)** *Left*: Color-coded myofibers according to CSA (μm$^2$) of Dhh cKO and ctrl 7 days post CTX injury. Scale bar: 250 μm. *Right*: Average CSA (μm$^2$) of Dhh cKO (n=10 TAs) and ctrl (n=6 TAs) mice 7 days post CTX injury. All data are represented as mean ± SEM. An unpaired two-tailed t test or a one-way ANOVA followed by a Dunnet's multiple comparison was used.

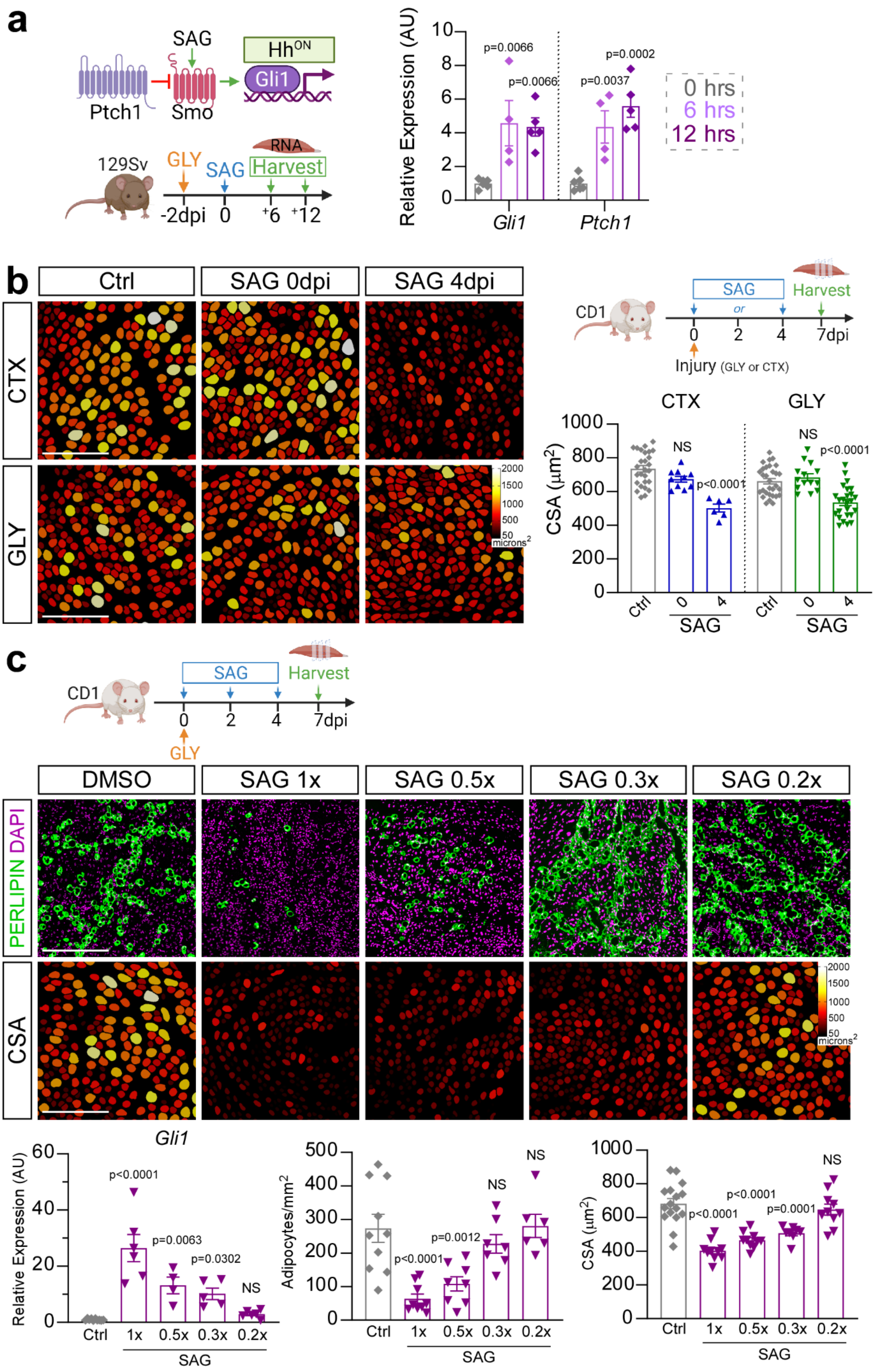

a
SAG
Hh^ON
Gli1
Ptch1
Smo
129Sv
GLY
SAG
RNA
Harvest
-2dpi
0
+6
+12
Relative Expression (AU)
p=0.0066
p=0.0066
p=0.0037
p=0.0002
Gli1
Ptch1
0 hrs
6 hrs
12 hrs
b
Ctrl
SAG 0dpi
SAG 4dpi
CTX
GLY
microns²
CD1
SAG
or
Harvest
7dpi
Injury (GLY or CTX)
CTX
GLY
CSA (μm²)
NS
p<0.0001
NS
p<0.0001
Ctrl
SAG
c
CD1
SAG
Harvest
7dpi
GLY
DMSO
SAG 1x
SAG 0.5x
SAG 0.3x
SAG 0.2x
PERLIPIN DAPI
CSA
microns²
Gli1
Relative Expression (AU)
p<0.0001
p=0.0063
p=0.0302
NS
Ctrl
1x
0.5x
0.3x
0.2x
SAG
Adipocytes/mm²
p<0.0001
p=0.0012
NS
NS
CSA (μm²)
p<0.0001
p<0.0001
p=0.0001
NS

**Supplemental Figure 3. SAG influences adipogenesis and myogenesis in a dose- and time-dependent manner.**

**a)** *Left:* Experimental outline. *Right*: After 2 days post GLY injury, RT-qPCR of *Gli1* and *Ptch1* at 0 hrs (n=6 TAs), 6 hrs (n=4 TAs) and 12 hrs (n=5 TAs) following SAG administration. **b)** *Left:* Myofibers color-coded according to size (μm$^2$) 7 days post CTX or GLY in ctrl and SAG- treated (at 0 dpi or 4 dpi) mice. Scale bar: 250 μm. *Right*: Average CSA (μm$^2$) 7 days post injury with vehicle control (CTX n=27 TAs & GLY n=30 TAs), SAG at 0 dpi (CTX: n=10 TAs & GLY: n=14 TAs) and SAG at 4 dpi (CTX: n=6 TAs & GLY: n=25 TAs). To note, these time points are part of the experiment described in the main Figure 3 and, thus, the same controls were used. **c)** SAG was administered at 0-, 2- and 4dpi at varying concentrations and TAs harvested at 7 days post GLY injury. Immunofluorescence of adipocytes (PERILIPIN$^+$, green) and nuclei visualized with DAPI (purple). Color-coded myofibers based on cross sectional area (CSA). Scale bars: 250 μm. *Bottom*: RT-qPCR of *Gli1* expression 7 days post GLY after treatment with DMSO control (n=10 TAs), SAG 1x (n=6 TAs), SAG 0.5x (n=4 TAs), SAG 0.3x (n=5 TAs) and SAG 0.2x (n=6 TAs). Quantification of adipocytes per injured area (mm$^2$) 7 days post GLY injury of DMSO control (n=10 TAs), SAG 1x (n=9 TAs), SAG 0.5x (n=8 TAs), SAG 0.3x (n=7 TAs), and SAG 0.2x (n=6 TAs). Average CSA (μm$^2$) 7 days after GLY injury of DMSO control (n=16 TAs), SAG 1x (n=10 TAs), SAG 0.5x (n=11 TAs), SAG 0.3x (n=8 TAs), and SAG 0.2x (n=10 TAs). All data are represented as mean ± SEM. One- way ANOVA followed by a Dunnet's multiple comparison was used.

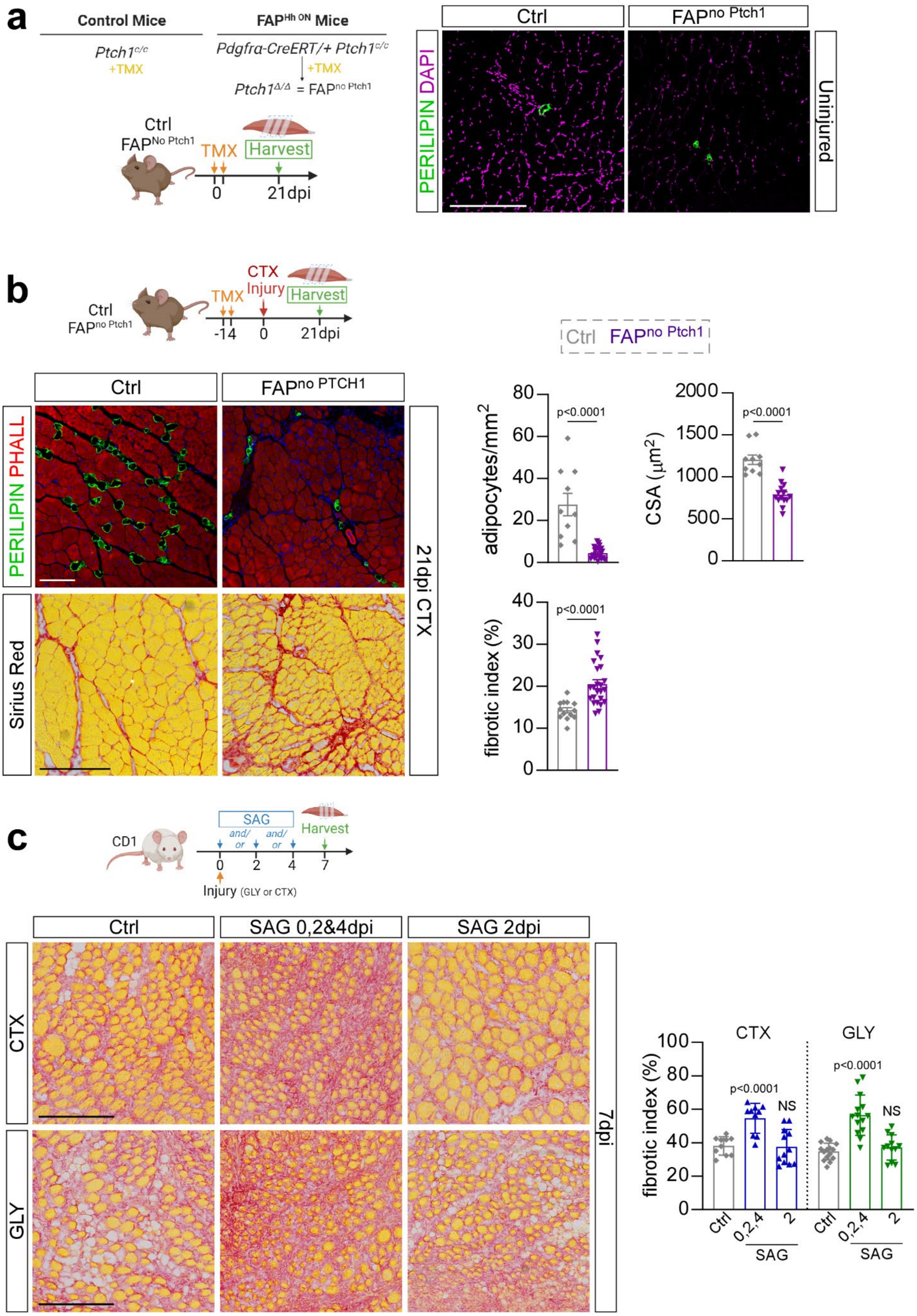

a
Control Mice
FAP^Hh ON Mice
Ptch1c/c
+TMX
Pdgfra-CreERT/+ Ptch1c/c
+TMX
Ptch1Δ/Δ = FAPno Ptch1
Ctrl
FAPNo Ptch1
TMX
Harvest
0
21dpi
Ctrl
FAPno Ptch1
PERILIPIN DAPI
Uninjured
b
Ctrl
FAPno Ptch1
TMX
CTX
Injury
Harvest
-14
0
21dpi
Ctrl FAPno Ptch1
Ctrl
FAPno PTCH1
PERILIPIN PHALL
Sirius Red
21dpi CTX
adipocytes/mm2
p<0.0001
CSA (μm2)
p<0.0001
fibrotic index (%)
p<0.0001
c
CD1
SAG
and/or
and/or
Harvest
0
2
4
7
Injury (GLY or CTX)
Ctrl
SAG 0,2&4dpi
SAG 2dpi
CTX
GLY
7dpi
CTX
GLY
fibrotic index (%)
p<0.0001
NS
p<0.0001
NS
Ctrl
0,2,4
2
SAG
Ctrl
0,2,4
2
SAG

**Supplemental Figure 4. Genetic and pharmacological Hh activation prevents IMAT formation but causes fibrosis and impairs myogenesis.**

**a)** *Left*: Experimental outline. *Right:* Immunofluorescence of PERILIPIN$^+$ adipocytes (green) of uninjured FAP$^{no\ Ptch1}$ and control mice (n=5 for each). Scale bars: 250 µm. **b)** Immunofluorescence of PERILIPIN$^+$ adipocytes (green) and PHALLOIDIN+ myofibers (scale bar: 100 µm) and histological Sirius red staining (scale bar: 250 µm) 21 days post CTX injury in FAP$^{no\ Ptch1}$ and control mice. *Right*: Average CSA (µm$^2$) 21 days post CTX of ctrl (n=10 TAs) and FAP$^{no\ Ptch1}$ (n=14 TAs) mice. Quantification of adipocytes per injured area (mm$^2$) 21 days post CTX injury of ctrl (n=10 TAs) and FAP$^{no\ Ptch1}$ (n=22 TAs) mice. Quantification of percent of collagen (red) of total TA area 21 dpi CTX in FAP$^{no\ Ptch1}$ (n=19 TAs) and ctrl (n=24 TAs). **c)** *Left:* Sirius red staining 7 days after CTX *(top)* or GLY *(bottom)* injury after vehicle control, SAG at 0-, 2- and 4 dpi, or SAG at 2 dpi. Scale bars: 250 µm. *Right:* Quantification of percent of Sirius red-positive collagen (red) over total TA area 7 days after injury from mice treated with vehicle control (CTX: n=10 TAs & GLY: n=15 TAs), SAG at 0-, 2- and 4 dpi (CTX: n=10 TAs & GLY: n=14 TAs) or SAG at 2 dpi (CTX & GLY: n=12 TAs). **c)** All data are represented as mean ± SEM. An unpaired two-tailed t test or a one-way ANOVA followed by a Dunnet's multiple comparison was used.

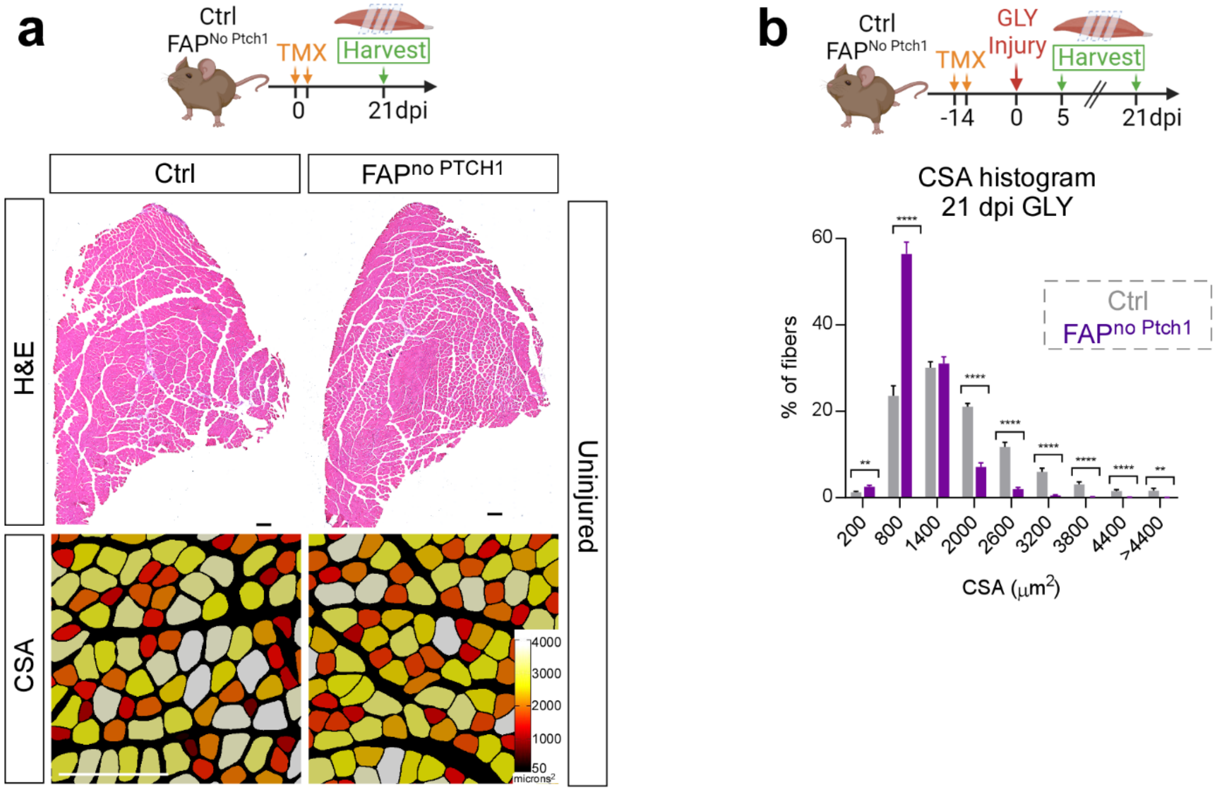


**Supplemental Figure 5. Ectopic Hh activation does not impact muscle homeostasis.**
**a)** Experimental outline. Color-coded myofibers based on cross sectional area (CSA) of uninjured FAP$^{no\ Ptch1}$ and control mice. Scale bar: 250 µm. H&E staining of uninjured FAP$^{no\ Ptch1}$ and control mice. Scale bar: 500 µm. **b)** *Top*: Experimental outline. *Middle*: Fiber number distribution, displayed as percent of total fibers, based on their CSA (µm$^2$) in FAP$^{no\ Ptch1}$ (n=18) and ctrl mice (n=14) 21 days post GLY injury. Source data are provided as a Source Data file. All data are represented as mean ± SEM. A two-way ANOVA followed by Tukey's multiple comparison test was used. A p value less than 0.05 was considered statistically significant where: ** p≤ 0.01, *** p≤ 0.001 and **** p≤ 0.0001. Source data are provided as a Source Data file.

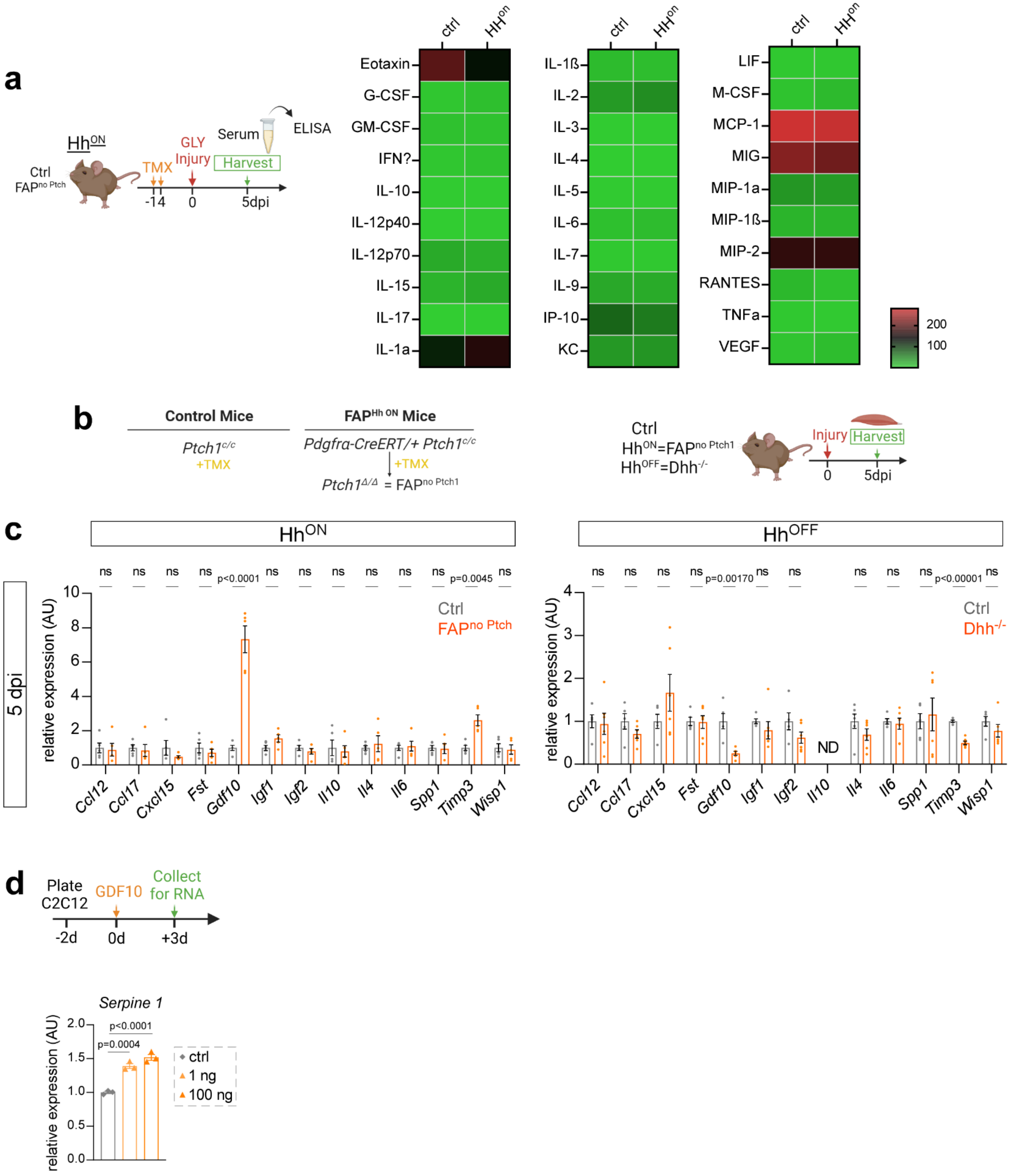

a
Hh^ON
Ctrl
FAP^no Ptch
TMX
GLY
Injury
Serum
Harvest
ELISA
-14
0
5dpi
ctrl
HH^on
Eotaxin
G-CSF
GM-CSF
IFN?
IL-10
IL-12p40
IL-12p70
IL-15
IL-17
IL-1a
IL-1ß
IL-2
IL-3
IL-4
IL-5
IL-6
IL-7
IL-9
IP-10
KC
LIF
M-CSF
MCP-1
MIG
MIP-1a
MIP-1ß
MIP-2
RANTES
TNFa
VEGF
200
100
b
Control Mice
FAP^Hh ON Mice
Ptch1^c/c
+TMX
Pdgfra-CreERT/+ Ptch1^c/c
+TMX
Ptch1^Δ/Δ = FAP^no Ptch1
Ctrl
Hh^ON=FAP^no Ptch1
Hh^OFF=Dhh^-/-
Injury
Harvest
0
5dpi
c
Hh^ON
Hh^OFF
5 dpi
relative expression (AU)
ns
p<0.0001
p=0.0045
Ctrl
FAP^no Ptch
p=0.00170
p<0.00001
Dhh^-/-
ND
Ccl12
Ccl17
Cxcl15
Fst
Gdf10
Igf1
Igf2
Il10
Il4
Il6
Spp1
Timp3
Wisp1
d
Plate C2C12
GDF10
Collect for RNA
-2d
0d
+3d
Serpine 1
p<0.0001
p=0.0004
ctrl
1 ng
100 ng

**Supplemental Figure 6. Screen for FAP-specific and Hh-induced factors that control adipogenesis and myogenesis.**
**a)** Enzyme-linked immunosorbent assay (ELISA) of whole muscle protein lysates from $FAP^{no\ Ptch1}$ and ctrl mice 5 days post GLY injury. **b)** Experimental outline. **c)** RT-qPCR of whole muscle RNA from $FAP^{no\ Ptch1}$ (n=5 TAs) and ctrls (n=5 TAs); and $Dhh^{-/-}$ (n=6 TAs) and ctrls (n=4-6) 5 days post injury for the known FAP targets: *Ccl2, Ccl7, Cxcl5, Fst, Gdf10, Igf1, Igf2, Il10, Il4, Spp1, Timp3* and *Wisp1*. **d)** RT-qPCR of *Serpine 1* from RNA isolated from C2C12 cells (n= 3 replicates per experimental group) 3 days after induction in ctrl versus rGDF10-treated cells (1 ng & 100 ng). All data are represented as mean ± SEM. A one-way ANOVA followed by a Dunnet's multiple comparison was used.

### Supplementary Table

#### Supplementary Table 1: List of primers used for RT-qPCR

| Gene | Forward | Reverse |
|---|---|---|
| *Hprt* | CATAACCTGGTTCATCATCGC | TCCTCCTCAGACCGCTTTT |
| *Pde12* | ACCTTTTGGGTGCCAGTAGA | CCAGAGGTCATCTGTCCTTCA |
| *Dhh* | CGATGGCTAGAGCGTTCAC | GTACCCAACTACAACCCCGA |
| *Timp3* | TAGACCAGAGTGCCAAAGGG | CCAGGATGCCTTCTGCAAC |
| *Gli1* | GGTGCTGCCTATAGCCAGTGTCCTC | GTGCCAATCCGGTGGAGTCAGACCC |
| *Ptch1* | AATTCTCGACTCACTCGTCCA | CTCCTCATATTTGGGGCCTT |
| *Gdf10* | CCAAATCCTTTGACGCCTACT | GCTCTGACGATGCTCTGGAT |
| Acta2 | GACAGAGGCACCACTGAACC | ACAGCACAGCCTGAATAGCC |
| *Ccl12* | TTAAAAACCTGGATCGGAACCAA | GCATTAGCTTCAGATTTACGGGT |
| *Ccl17* | GCTGCTTTCAGCATCCAAGTG | CCAGGGACACCGACTACTG |
| *Cxcl15* | GTTCCATCTCGCCATTCATGC | GCGGCTATGACTGAGGAAGG |
| *Fst* | GCCTCCTGCTGCTGCTACTC | TTATACAGGACCTGGCAGCG |
| *Igf1* | CTGAGCTGGTGGATGCTCTT | TCATCCACAATGCCTGTCTG |
| *Igf2* | GTGCTGCATCGCTGCTTAC | ACGTCCCTCTCGGACTTGG |
| *Il10* | AGCATTTGAATTCCCTGGGT | TTTTCACAGGGGAGAAATCG |
| *Il4* | GGTCTCAACCCCCAGCTAGT | GCCGATGATCTCTCTCAAGTGAT |
| *Il6* | TAGTCCTTCCTACCCCAATTTCC | TTGGTCCTTAGCCACTCCTTC |
| *Spp1* | AGCAAGAAACTCTTCCAAGCAA | GTGAGATTCGTCAGATTCATCCG |
| *Wisp1* | GTGTGATGATGACGCAAGGA | ATGCAGTTCTCATACCGTTGC |
| *Serpine 1* | GTGAATGCCCTCTACTTCAGTG | GCTGCCATCAGACTTGTGGAA |
